# Werewolf, there wolf: variants in *Hairless* associated with hypotrichia and roaning in the lykoi cat breed

**DOI:** 10.1101/2020.05.07.082719

**Authors:** Reuben M. Buckley, Barbara Gandolfi, Erica K. Creighton, Connor A. Pyne, Michelle L. LeRoy, David A. Senter, Delia M. Bouhan, Johnny R. Gobble, Marie Abitbol, Leslie A. Lyons, 99 Lives Consortium

**Affiliations:** Department of Veterinary Medicine and Surgery, College of Veterinary Medicine, University of Missouri, Columbia, MO, USA; Veterinary Allergy and Dermatology Clinic, LLC., Overland Park, KS 66210 USA; Tellico Bay Animal Hospital, Vonore, TN 37885 USA; NeuroMyoGène Institute, CNRS UMR5310, INSERM U1217, Faculty of Medicine, Rockefeller, Claude Bernard Lyon I University, Lyon, France; Univ Lyon, VetAgro Sup, Marcy-l’Etoile, France

**Keywords:** atrichia, domestic cat, *Felis catus*, fur, *HR*, naked

## Abstract

A variety of cat breeds have been developed via novelty selection on aesthetic, dermatological traits, such as coat colors and fur types. A recently developed breed, the lykoi, was bred from cats with a sparse hair coat with roaning, implying full color and all white hairs. The lykoi phenotype is a form of hypotrichia, presenting as significant reduction in the average numbers of follicles per hair follicle group as compared to domestic shorthair cats, a mild to severe perifollicular to mural lymphocytic infiltration in 77% of observed hair follicle groups, and the follicles are often miniaturized, dilated, and dysplastic. Whole genome sequencing was conducted on a single lykoi cat that was a cross between two independently ascertained lineages. Comparison to the 99 Lives dataset of 194 non-lykoi cats suggested two variants in the cat homolog for *Hairless* (*HR*: *lysine demethylase and nuclear receptor corepressor*) as candidate causal variants. The lykoi cat was a compound heterozygote for two loss of function variants in *HR*, an exon 3 c.1255_1256dupGT (chrB1:36040783), which should produce a stop codon at amino acid 420 (p.Gln420Serfs*100) and, an exon 18 c.3389insGACA (chrB1:36051555), which should produce a stop codon at amino acid position 1130 (p.Ser1130Argfs*29). Ascertainment of 14 additional cats from founder lineages from Canada, France and different areas of the USA identified four additional loss of function *HR* variants likely causing the highly similar phenotypic hair coat across the diverse cats. The novel variants in *HR* for cat hypotrichia can now be established between minor differences in the phenotypic presentations.

## 1. Introduction

Domestic cats have been developed into distinctive breeds during the past approximately 150 years, since the first cat shows held in the late 1800’s [1–3]. Many breeds have proven to be genetically distinct [4,5] but also suffer from inbreeding and founder effects, inadvertently becoming important biomedical models for human diseases. Over 72 diseases / traits caused by at least 115 mutations have been discovered in cat breeds (https://omia.org/) [6,7]. To produce novel breeds, cats have been selected mainly for aesthetic, dermatological traits since the phenotypes can be easily recognized by cat enthusiasts, the unique appearance leading to a new breeding program. A majority of breeds were developed after the World Wars and several are defined by interesting coat DNA variants, such as the Cornish rex [8], Devon rex, sphynx [9], and the Selkirk rex [10,11]. These coat mutations are innocuous in the cat, but the same genes for atrichia and hypotrichia cause ectodermal dysplasias in humans [12–15] and other species [16–22]. However, some cat coat and fur types are associated with maladies. The *FOXN1* variant that causes a hypotrichosis in cats is associated with a health condition and shortened life expectancy in the Birman breed [23]. The *White* locus variant in *KIT* has pleiotrophic effects in ocular tissues and is associated with deafness [24]. Albinism and temperature-sensitive variants in *tyrosinase* (*TYR*) [25,26], the *Color* locus in cats, are associated with disruption of the optical chiasma, leading to strabismus and nystagmus [27]. But overall, a majority of cat fur types and coat colors have few detrimental health effects.

A recently developed breed of cat, termed the lykoi (**Figure 1**), presents a unique form of hypotrichia [28]. Lykoi have a significant reduction in the average numbers of follicles per hair follicle group as compared to domestic shorthair cats, a mild to severe perifollicular to mural lymphocytic infiltration in 77% of observed hair follicle groups, and the follicles are often miniaturized, dilated, and dysplastic. Individual hairs of the coat are either normal coloration or all white, producing a roaning effect. The undercoats are sparse. The lykoi has been genotyped for all the known cat fur type mutations, including variants in *KRT71*, which cause the hairless sphynx breed, Devon rex [9] and Selkirk rex [11] curly hair, and none of these variants are present in the lykoi cats. The breeding program was established in 2011 by a veterinarian, who has constantly monitored health in the cats [29]. No health concerns have been identified in the lykoi other than the lymphocytic mural folliculitis.

**Figure 1.**
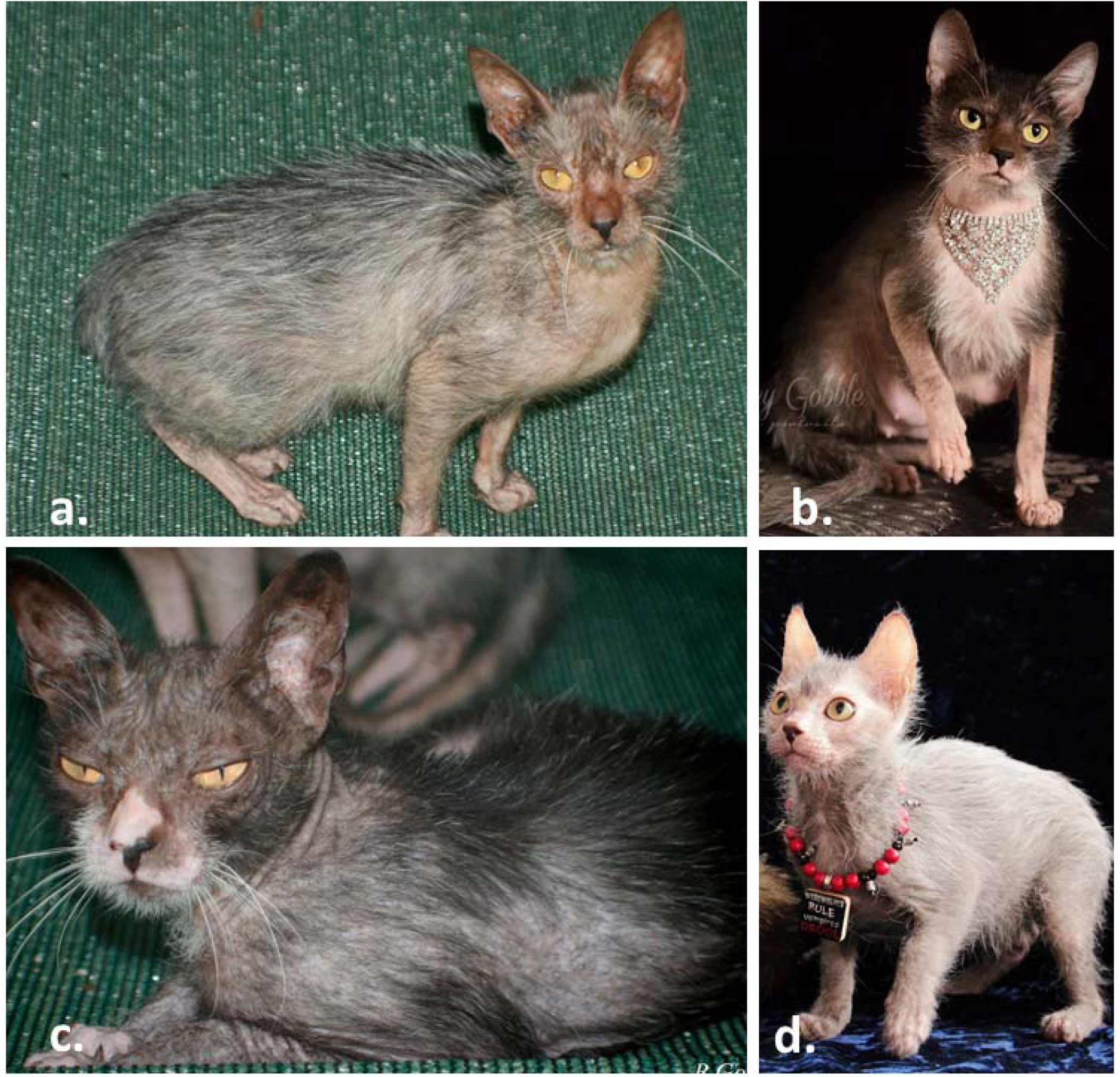
The lykoi cat breed. Lykoi breed founders from independent sightings identified in 2010. a.) Virginia lineage (c.3389insGACA), b.) Missouri lineage (c.1255_1256 dupGT), c.) Tennessee lineage (c.1255_1256dupGT), d.) Canadian lineage (c.2593C>T). Solid black is the preferred coloration as the roaning of the white hairs is more distinctive. Note sparse hair on the lower limbs. (Images courtesy of Brittney Gobble). Sixteen different lineages were ascertained containing six different variants in *HR*. Cats can molt their hair coat at different times during development and through-out the year.

Whole genome sequencing (WGS) has proven a successful genetic approach for the identification of causal variants for several phenotypes and diseases in the domestic cat [30–33]. This study used WGS to identify the causal variant(s) for the lykoi presentation in the domestic cats.

## 2. Materials and methods

### 2.1. Ethics statement

All procedures performed in studies involving animals were in accordance with the ethical standards of the University of Missouri (MU) institutional animal care and use protocol 8701 and 8313. All samples were collected with informed owner consent.

### 2.2 Lykoi samples

Samples for DNA isolation from the lykoi cats were provided voluntarily with the permission of the owners as either whole blood EDTA or buccal swabs. DNA was isolated by organic methods [34] or using DNAeasy kits (Qiagen, Valencia, CA) according to the manufacturer’s protocol. To develop pedigrees, the breeder/owner reported parentage of submitted cats, parentage was verified with a panel of feline-derived short tandem repeats (STRs) as previously described [35]. STR fragment sizes were determined using STRand analysis software [36]. Samples from unrelated cats with similar phenotypes were also ascertained (**Table 1**).

**Table 1.**
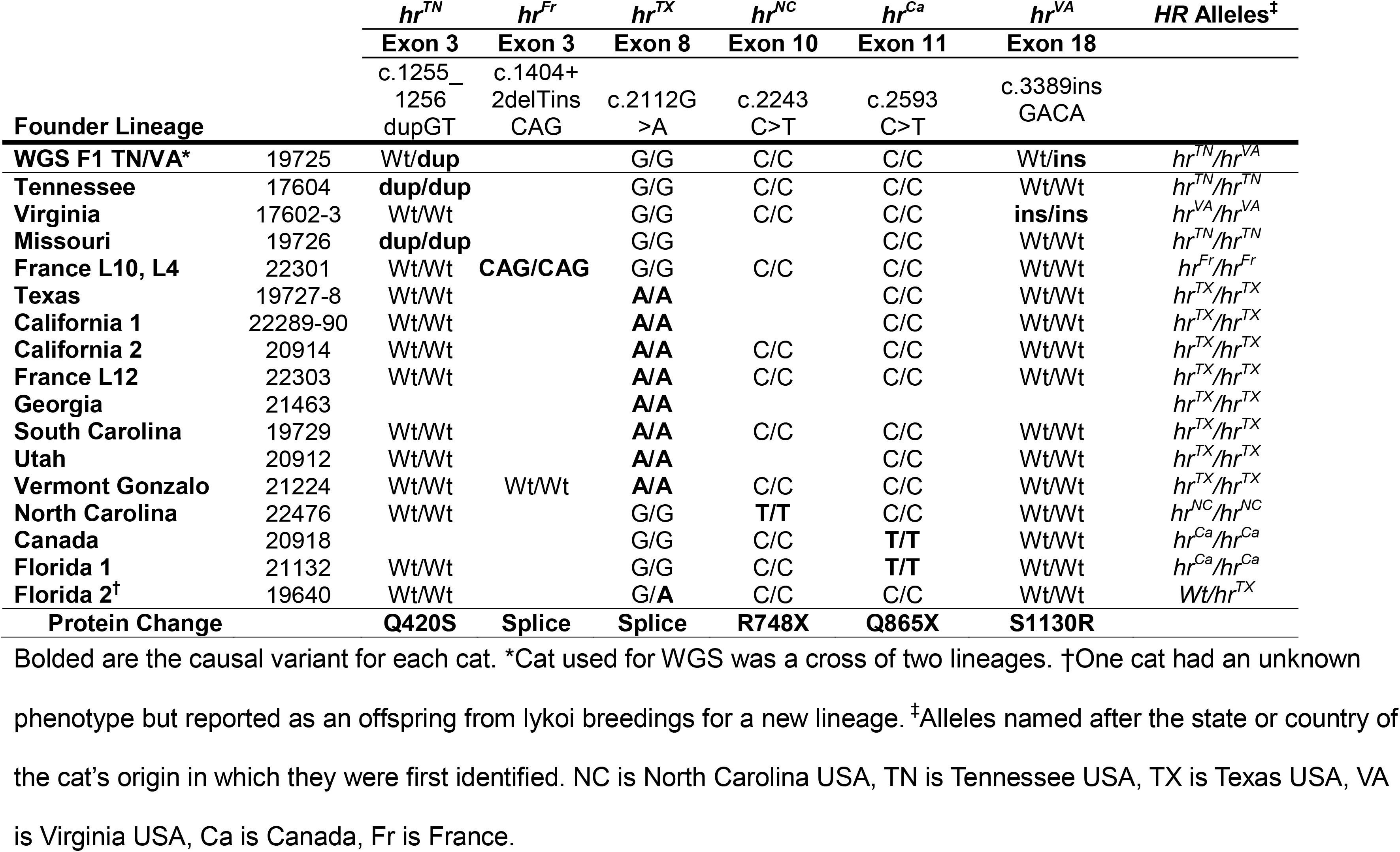
Lykoi founder lineage variants in *Hairless* (*HR*) for cats with the hypotrichia presentation.

### 2.3 Whole genome sequencing and variant calling

A single lykoi cat was whole genome sequenced as previously described [32]. The selected cat was an F1 from the mating of two independently discovered foundation lineages from Virginia and Tennessee. The sequence was included in the 195 - cat analysis of the 99 Lives cat genome sequencing project and submitted to the NCBI short read archive under BioProject: PRJNA308208, PRJNA288177; BioSample: SAMN05980355. For the 195 - cat analysis, reads were mapped to Felis_catus_9.0 [37] and assigned to read groups using BWA-MEM from Burrows-Wheeler Aligner version 0.7.17 [38]. Duplicate reads were marked using MarkDuplicates from Picard tools version 2.1.1 (http://broadinstitute.github.io/picard/), with OPTICAL_DUPLICATE_PIXEL_DISTANCE set at 2500. Genome Analysis Toolkit version 3.8 (GATK 3.8) was used to further process the sequence data [39]. Indel realignment was performed with RealignerTargetCreator and IndelRealigner [39] and SNPs, and Indels were called using HaplotypeCaller in gVCF mode (-ERC GVCF) [40]. The gVCFs were combined into groups of ∼20 individuals using CombineGVCFs and were genotyped simultaneously using GenotypeGVCFs. Throughout, Samtools version 1.7 sort, index, view, and cat functions were used to process BAM files between individual tasks [41]. Together these processes produced a single VCF comprised of 195 cats for downstream analysis. Code used to process individual genomes is publicly available on github (https://github.com/mu-feline-genome/github-lewis/blob/master/map_libraries.slurm.sh). DNA variants were viewed, filtered and annotated using VarSeq (Golden Helix, Boseman, MT) with the Ensembl release 98 Felis_catus_9.0 genome annotation [42]. Candidate variants were considered to be homozygous or compound heterozygous in the same gene in the lykoi cat and not present in any other cat of the 99 Lives cat database. Only variants that caused high to moderate severity effects on the protein were considered and variants with high severity and within candidate genes were prioritized. Sequencing primers were developed for candidate variants as previously described [9] for the homolog of *HR* using sequences NCBI Accessions: XM_023252512.1, XM_011281452.3 (**Supplementary Table 1**).

### 2.4 Hairless (HR) genotyping and sequencing

The two *HR* frameshift variants, including an exon 3 c.1255_1256dupGT, and the exon 18 c.3389insGACA, were identified by the WGS analyses. These variants were validated in the WGS cat by Sanger sequencing (**Supplementary Table 1**). An assay was designed as previously described [32] to genotype the identified variants in pedigree A (**Supplementary Figure 1**) and the additional cats, using the Agena Bioscience iPLEX Gold Genotyping reagent set (Agena Bioscience Inc., San Diego, CA) (**Supplementary Table 2**). Products were genotyped with the MassARRAY System with Nanodispenser RS1000 (Agena Bioscience Inc., San Diego, CA).

Not all ascertained cats with similar hair coats had the WGS identified variants, therefore the coding regions of *HR* were Sanger sequenced in each additional founder cat (**Supplementary Table 1**). PCR and thermocycling conditions were conducted as previously described [43]. The variants for the cats in the pedigree B (**Supplementary Figure 2**) were also genotyped by Sanger sequencing.

## 3. Results

### 3.1 Lykoi samples

Over 100 cats were ascertained for the lykoi project and were used to develop two pedigrees of the cats segregating for the lykoi phenotype (**Supplementary Figures 1 and 2**). The relationship of the cats was confirmed by STRs (data not shown). Sixty-seven cats formed an extended pedigree “A” by crossing three different lineages (Tennessee, Virginia and Texas) (**Supplementary Figure 1**) and a smaller pedigree “B” was obtained from a French lineage of cats (**Supplementary Figure 2**). Overall, cats were identified from 16 foundation lines, ascertained from 14 diverse regions in the USA, Canada and France. Two supposed founder lineages were independently ascertained from Florida, California and France, each (**Table 1, Figure 1**).

### 3.2 Whole genome sequencing

The selected cat for the WGS represented two founder lineages (**Supplementary Figure 1**) and a mean of 48.4x genomic sequence coverage was produced for the sequenced cat. Approximately 558 variants were identified as heterozygous in the lykoi cat. Seventeen were loss of function variants and 154 were missense variants (**Table 2, Supplementary File 1**). Only one gene was identified with variants that caused highly severe effects on the protein. The two variants in the cat homolog of *Hairless* (*HR*), *lysine demethylase and nuclear receptor corepressor* (cat chromosome B1:36,038,754 – 36,052,521), were considered the highest priorities as both variants have severe effects and supported the suspected compound heterozygosity in the sequenced lykoi cat. Additionally, *HR* is a known gene causing atrichia in mice [44] and humans [45]. The lykoi cat was a compound heterozygote for two loss of function variants in *HR* transcript (HR-202 ENSFCAT00000012982.5); specifically, an exon 3 c.1255_1256dupGT (chrB1:36040784), which should produce a stop codon at amino acid 420 (p.Gln420Serfs*100) in the Tennessee lineage and is designated *hr^TN^* allele and, an exon 18 c.3389insGACA (chrB1:36051556), which should produce a stop codon at amino acid position 1130 (p.Ser1130Argfs*29) in the Virginia lineage and is designated *hr^VA^* allele (**Figure 2**). These two identified frameshift variants were confirmed by direct Sanger sequencing in the cat submitted for WGS and the presented positions are for the newest cat genome assembly Felis_Catus_9.0 (GCF_000181335.3/). The lykoi phenotype segregated concordantly with each loss of function variant across the pedigree developed from the Virginia and Tennessee lineages (**Supplementary Figure 1**). Cats with the lykoi hair coat in these lineages were either homozygous for one of the two loss of function variants or compound heterozygous for both loss of function variants.

**Figure 2.**
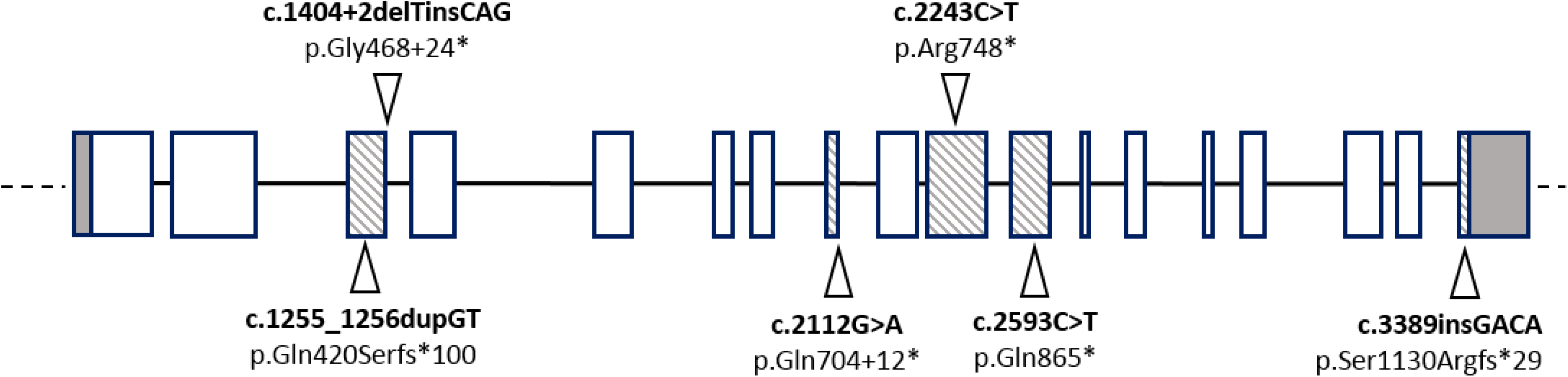
*HR* gene and variants in the lykoi breed. Genomic location of the identified variants associated with the *Hairless* phenotype in the lykoi breed. UTR regions are presented in grey while exons that contain one of the six identified variants are shaded. Variants location are identified by triangles, two of the identified variants disrupt a splicing site (c.1404+2delTinsCAG and c.2112G>A) while all other variants (c.1255_1256 dupGT, c.,2243C>T, c.,2593C>T and c.3389insCAGA) are predicted to produce a truncated protein product.

**Table 2.** Unique 99 Lives heterozygous WGS variants in a lykoi cat.

| Severity | Effect | No. | Genes | HR |
| --- | --- | --- | --- | --- |
| High | Frame Shift (LOF) | 13 | 12* | 2 |
|  | Splice donor - acceptor | 0 | 0 | 0 |
|  | Stop gained (LOF) | 4 | 5 | 0 |
| Moderate | Missense | 154 | 146 | 62 |
| Low | Splice region | 31 | 30 | 5 |
|  | Synonymous | 127 | 120<sup>†</sup> | 67 |
| Other | **Intronic** | 118 | 103 | 9 |
|  | **Intergenic & UTR** | 17 | -- | 7 |
|  | **Non-coding exon** | 94 | 68 | 0 |
|  | **Total variants** | 558 | ~426** | 152 |

\*Two variants in *HR*, <sup>†</sup>including p.Ser1130Arg in *HR*. Three variants in *HR* were unique to the lykoi cat in the WGS comparison to 193 additional cats. \*\*Total includes undefined transcripts and suspected coding sequences.

WGS data also revealed additional *HR* variants. There was one synonymous variant (p.Val1129=), two missense variants (p.Lys433Asn and p.Ser1130Arg), 14 intronic variants, and one 3’ UTR variant. One of the missense variants was an exon 3 c.1299A>C, suggesting a p.Lys433Asn amino acid change of a positively charged lysine to a polar and uncharged asparagine. The p.Lys433Asn was a common variant with an allele frequency of 0.57. Conversely, the p.Ser1130Arg variant was heterozygous in only one other cat in the 99 Lives dataset (**Table 2, Supplementary File 2**). Heterozygous only splice region variants were identified in the 99 Lives dataset that appeared to be derived from Bengal cats, hence perhaps of Asian Leopard cat (*Prionailurus bengalensis*) origin.

### 3.3 Lykoi variants in other lineages

The *HR* variants discovered using WGS (c.1255_1256dupGT and c.3389insGACA) were absent from other lykoi cats from different lineages, suggesting multiple causative variants for the phenotype. To identify additional lykoi variants, direct sequencing of the coding region of *HR* was performed on lineage founders. Four additional variants were identified (**Figure 2**, **Table 1**). Firstly, an exon 3 splice variant c.1404+2delTinsGT (chrB1:36040933) was identified in a cat from France Pedigree B and is designated *hr^Fr^* allele (**Figure 2, Supplementary Figure 2**). This variant should extend and change the reading frame, including an additional 24 amino acids in the aberrant protein before a stop codon is recognized. Alternatively, a cryptic splice site may be used from within intron 4. Secondly, an exon 8 variant at c.2112G>A (chrB1:36045776) was identified in seven different lineages as homozygous, including a cat from France and is heterozygous in a suspected obligate carrier from Florida. This variant likely disrupts the splice donor allowing read through for an additional 12 amino acids until a stop codon is encountered and is designated *hr^TX^* allele. Alternatively, a cryptic splice site may be used from within intron 8. Finally, two additional stop codon producing variants were also identified including an exon 10 c.2243C>T (p.Arg748X) (chrB1:36047047) in a cat from North Carolina, designated *hr^NC^* allele, and an exon 11 c.2593C>T (p. Gln865X) (chrB1:36047518) identified in two cats from Tennessee and Canada and is designated *hr^Ca^* allele. Five submitted founder cats had unique variants. The founder cat from Canada had the same variant as one of the submitted founders from Florida. The other cat from Florida had a normal coat but was the offspring of a suspected new lineage. This cat was heterozygous for the exon 8 splice site variant thus, the queen did not have a novel variant. Overall, six likely causal variants were identified (**Figure 2**) in 16 lineages, including seven lineages from the USA covering 11 states and one cat from France sharing the same exon 8 splice site variant. The variants, positions and flanking sequences are presented in **Supplementary File 3**.

Known crosses of the different foundation lineages supported the causal function of the identified variants. Sixteen cross lineage cats that had the lykoi hair coat were compound heterozygotes, including 16 for the *hr^TN^/hr^VA^ alleles* (**Supplementary Figure 1**) and compound heterozygous lykoi cats with the *hr^TX^/hr^VA^* alleles in both pedigrees (**Supplementary Figure 1, 2**). One of the seven cats with the exon 3 c.1299A>C non-synonymous variant was also homozygous for the exon 10 c.2243C>T (p.Arg748X) stop codon variant and one other cat was homozygous for the exon 8 splice variant, further suggesting this missense variant as non-causal. The cats from France had been cross bred with cats from Italy and the USA, demonstrating the presence of the exon 8 and exon 18 variants. The French pedigree (Pedigree B – **Supplementary Figure 2**) also segregated for a novel exon 3 splice variant, indicating a novel *de novo* variant from Europe.

## 4. Discussion

Although over 50 cat breeds are identified by different cat associations and registries worldwide, fewer than 30 are demonstrated to be genetically distinct [4,5,46]. Novel cat breeds are continually being developed by producing a new breed from crosses with existing breeds, such as the Ocicat and Burmilla, by interbreeding domestic cats with small wild felids, such as Bengals and Savannahs, and by identifying new phenotypic variants in feral populations i.e., novelty selection, such as Devon [47], Cornish [48] and Selkirk rex [10]. Novelty breeds, such as Selkirk rex and Scottish folds, are characterized by novel “breed-defining” variants, retain high genetic variation [4,5,10], but often modify their type but by cross - breeding with established breeds that have the desired structural “look”. For example, the Selkirk rex has strong genetic influences from Persians and British shorthair [10], although the curly coat is a novelty phenotype identified in the past few decades in Northwestern USA [10].

The lykoi is a very recently developed novelty breed with a sparse hair coat and black and white hair roaning, hence named from the Greek term *lycos* for wolf. To maintain diversity in the founding population, the breeders have actively recruited cats with similar phenotypes for the breeding program, resulting in six different “foundation” lineages identified in this study from 16 potential founders. The breed is growing in popularity due to the novelty of the appearance, the lack of concern for health problems and the charismatic name and nature. The breed was accepted for full championship showing by TICA in May 2017 [29].

*Hairless* (*Hr*) (a.k.a. *lysine demethylase and nuclear receptor corepressor*) is one of the earliest mutations identified in mice (MMu Chr14:70554056-70573548) and over 30 phenotypic mutations have been identified, including ∼ 17 that are spontaneous, naturally occurring (MGD) [49]. The hairless mouse [50] is an insertion of murine leukemia proviral sequences into intron 6 resulting in aberrant splicing [51]. The *HR* gene encodes a protein that is involved in hair growth. This protein functions as a transcriptional corepressor of multiple nuclear receptors, including thyroid hormone receptor [52], the retinoic acid receptor-related orphan receptors [53] and the vitamin D receptors [54], and also interacts with histone deacetylases [55]. By modulating the activity of receptors, *HR* plays a critical role in skin function and hair maintenance by regulating both gene expression as well as epithelial stem cells differentiation. The translation of this protein is modulated by a regulatory ORF that exists upstream of the primary ORF, hence, the protein expression regulation is an overall critical element in directing hair growth [56]. The human homolog, *HR*, is on human chromosome 8p21.3; chr8:22114419-22131053. ClinVar lists 187 variants involving *HR*, 117 are limited to the gene and 17 are pathogenic or likely pathogenic mutations in humans [57]. Several *HR* variants are known to cause abnormalities in humans, such as alopecia universalis congenita (OMIM:203655) [52], atrichia with papular lesions (OMIM:209500) [58], which is an alopecia characterized by irreversible hair loss during the neonatal period on all hair-bearing areas of the body followed by the development of papular lesions, and Hypotrichosis 4, (a.k.a.) Marie Unna Type, 1; (OMIM:146550), which is caused by autosomal dominant mutations in the upstream ORF – U2RH [56]. Variants in *HR* in other species are relatively rare, but causal variants of hairless are known in sheep [17], atrichia with papular lesions is also identified in macaques [16], and, in dolphins, evolutionary loss has led to *HR* as a pseudogene, leading to hypotrichosis in this mammal [20]. Various other genes cause the hairless phenotypes, such as, *KRT71* in the sphynx breed [9], and *FOXI3* [19] and *SGK3* [59,60] in dogs.

*HR* in the cat is annotated in Ensemble 98 [61] as ENSFCAG00000012978 B1:36034352-36051895:1. Three transcripts are described containing 17 – 19 exons, in which exons 17 – 19 are the variable exons. Three 5’ UTRs are recognized, one as part of the 5’ portion of exon 1. Two transcripts have short 3’UTRs at the end of exon 18. The variants in this study were annotated with Ensembl 98 transcript HR-202 containing 4227 bp that translate to 1184 amino acids. Each of the six variants identified in the lykoi cats either cause termination codons at the variant site or cause downstream terminations after an additional 12 (exon 8 c.2112G>A) to 100 (exon 3 c.1255_1256dupGT) amino acids, leading to proteins with ∼528 – 704+12 amino acids. Interestingly, one variant, exon 18 c.3389insGACA (p.Ser1130Argfs*29), while associated with the phenotype, produces an almost full length protein (95%), suggesting the terminal end of the protein is required for normal function.

Several phenotypic traits in cats are heterogeneous, including the variants for the loci *Long*, *Tailless*, and the classic (blotched) pattern of *Tabby*, which are each caused by four different mutations in the genes *FGF5* [62,63], *TBX1* [64], and *LVRN* [65], respectively. Variation in the phenotypic presentations caused by these different variants is undocumented. Only the *TBX1* variants define breeds, the Manx and Cymric, which is a longhaired Manx, the *Long* and *Tabby* variants segregate within and amongst breeds. A few breeds have unique and breed defining variants, such as Scottish folds [66], Selkirk rex [11], Devon rex, and sphynx [9]. Like the Manx, the lykoi will be a unique breed that segregates for several variants within the same gene, *HR*, that present a similar phenotype (**Figure 1**). Unlike the Manx variants [64], the variants that cause the hypotrichosis are recessive and do not cause additional health concerns known to date. The only documented abnormality is the sparse haircoat resulting from abnormal follicular development and lymphocytic mural folliculitis. Some variants are not perpetuated as they tend to cause more periodic hair loss, suspected to be associated with sex hormone levels (JRG, personal communication). The lykoi breeders can now use genetic testing to monitor the variants in the population and to realize possible associations with phenotypic differences in compound heterozygotes. Additional haplotype analyses of flanking variants could determine if the eight reported founder lineages with the exon 8 variant are identical by descent or identical by state and represent multiple *de novo* mutation events at the same site.

## Supporting information

suppl_F3_lykoi_HR_sequences

suppl_F1_Lykoi_all_variants

suppl_F2_HR_variants

## Acknowledgements

Funding was provided in part by the Gilbreath McLorn Endowment of the College of Veterinary Medicine, University of Missouri, Cat Health Network (D14FE-552), the Winn Feline Foundation (W16-030) (LAL). We appreciate the assistance of cat breeders, including Patti Thomas, Cheryl Kerr and Christine Boulanger. Photographs were provided courtesy of Brittney Gobble. We appreciate technical assistance and support with the manuscript from Thomas R. Juba, technical assistance from Nicholas A. Gustafson, and assistance with figures from Karen Clifford.

## Authors’ contributions

Reuben M. Buckley, Barbara Gandolfi, Erica K. Creighton, Connor A. Pyne, Michelle L. LeRoy, David A. Senter, Delia M. Bouhan, Johnny R. Gobble, Marie Abitbol, Leslie A. Lyons^1^

- Conception and design – LAL, JRG, BG
- Provision of study materials – LAL, JRG, BG, MLL, DAS, MA
- Collection and assembly of data – LAL, EKC, CAP, JRG, DMB
- Analysis and interpretation of the data – LAL, BG, RMB, MA
- Drafting of the article – LAL, BG
- Obtaining of funding – LAL
- Critical revision of the article for important intellectual content – LAL, BG, RMB, EKC JRG, MA
- Final approval of the article – all authors

## Compliance with ethical standards

### Ethical approval

All procedures performed in studies involving animals were in accordance with the ethical standards of the University of Missouri institutional animal care and use protocol 8701 and 8313.

---

**Supplementary Figure 1.**
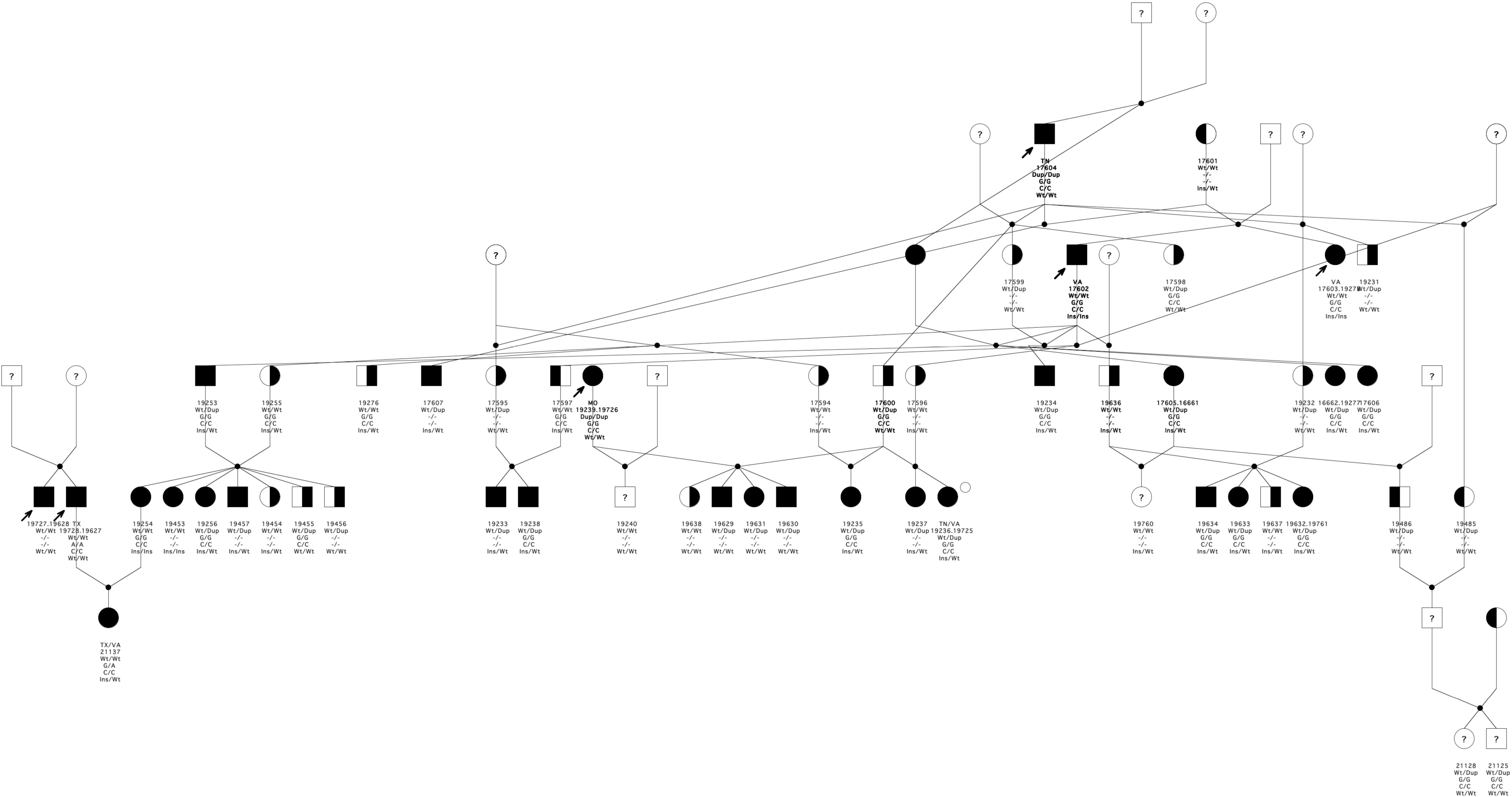
USA lineages of lykoi cats – Pedigree A. Virginia and Tennessee lines were initially crossed to develop the lykoi breed. Additional cats, from Missouri and Canada, were subsequently added to the pedigree and used in the breeding program. Relationships of 67 cats provided by the breeder and confirmed with genetic testing of STRs when possible (data not shown). Arrow indicates the probands of each lineage. Circles indicate females, squares indicate males, and diamonds indicate unknown sex. Filled symbols represent cats with the lykoi hair coat. Half-filled represent obligate carriers. Symbols with question marks represent cats with unknown phenotype. A symbol with no fill indicates the cat is known to be completely unrelated and not expected to be a carrier. Cats are identified by a laboratory number. An open circle at the upper right of symbol is the cat that was whole genome sequenced. Genotype of the two *HR* frameshift variants, the exon 3 c.1255_1256dupGT (*hr^TN^*), the exon 8 c.2112G>A stop codon variant (*hr^TX^*), the exon 11 c.2593 C>T (*hr^Ca^*), and the exon 18 c.3389insGACA (*hr^VA^*) is listed below the symbol.

**Supplementary Figure 2.**
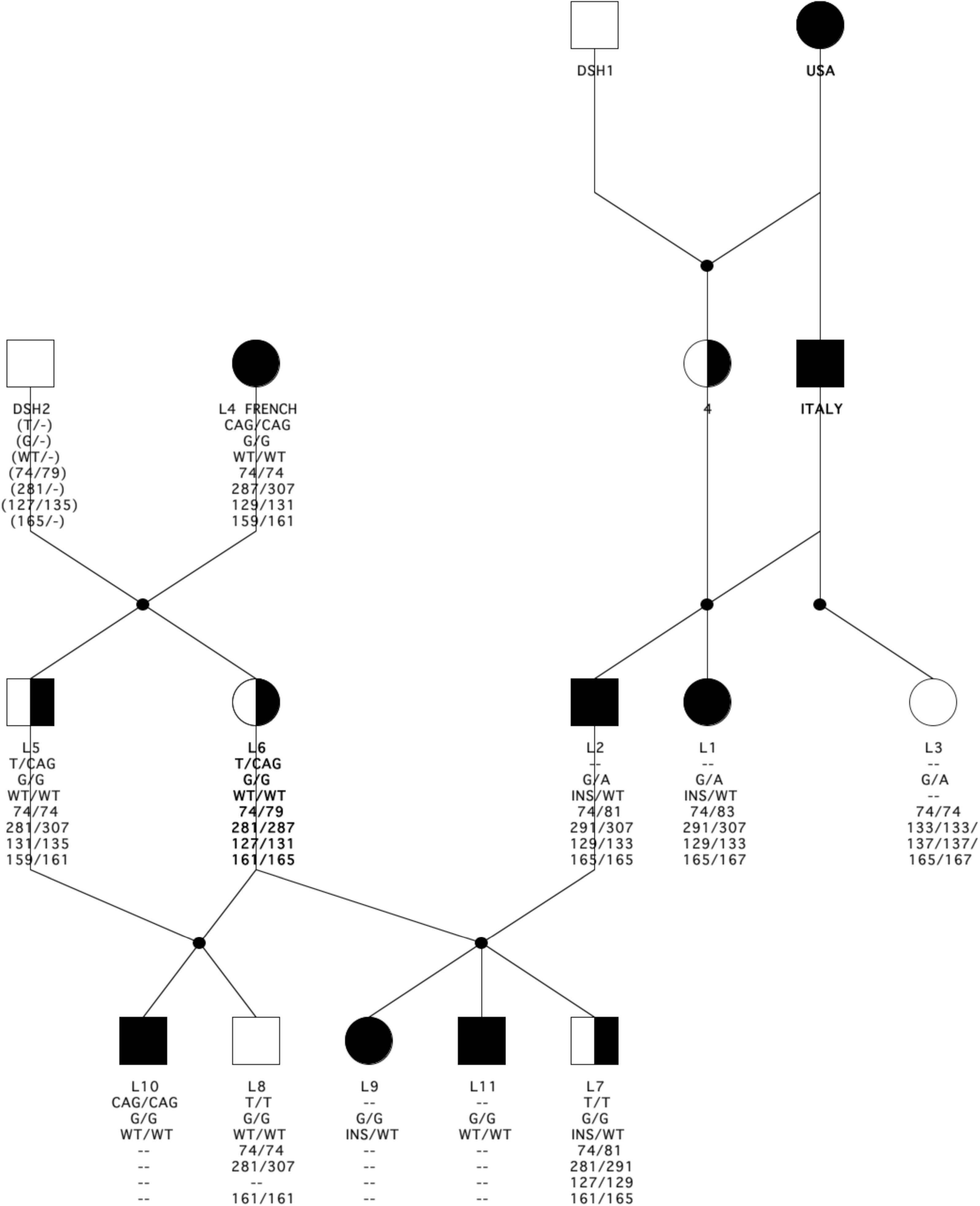
French lineages of lykoi cats - Pedigree B. Cats were identified in France with the lykoi phenotype. Relationships of 16 cats provided by the breeder and confirmed with genetic testing of STRs when possible. Arrow indicates the probands of each lineage. Circles indicate females, squares indicate males. Filled symbols represent cats with the lykoi hair coat. Half-filled represent obligate carriers with normal hair coat. A symbol with no fill indicates the cat is known to be completely unrelated and not expected to be a carrier. Presented under each symbol are the genotypes in the *HR* exon 3 c.1402+2T>CAG splice variant (*hr^Fr^*), the exon 8 c.2112G>A stop codon variant (*hr^TX^*) and the exon 18 insertion c.3393insGACA variant (*hr^VA^*). WT implies wildtype and INS implies insertion. Below the *HR* genotypes are the allelic sizes in basepairs for STRs: FCA149, F85, FCA075, and FCA229 [67,68]. Inferred genotypes are presented in parentheses. Cats L1 and L2 are compound heterozygote cats for variants previously identified in cats from the USA and Italy. Cat L3 has the exon 8 variant, likely originating from the USA foundation queen and inherited by L1 and L2 via cat 4. Cat L5 has the novel exon 3 c.1402+2T>CAG splice variant, inherited from the French founder cat L4. L6 produced five kittens from sires L2 and L5.

## References

1. Crystal Palace - Summer concert today Cat Show on July 13. Penny Illustrated Paper, Amusement: 1871, 510, July 08, 11.

2. The First Cat Show in America. New York Times March 06, 1881.

3. Morris, D. Cat breeds of the world; Penguin Books: New York, 1999.

4. Lipinski, M.J.; Froenicke, L.; Baysac, K.C.; Billings, N.C.; Leutenegger, C.M.; Levy, A.M.; Longeri, M.; Niini, T.; Ozpinar, H.; Slater, M.R., et al. The ascent of cat breeds: genetic evaluations of breeds and worldwide random-bred populations. Genomics 2008, 91, 12–21, doi:10.1016/j.ygeno.2007.10.009.

5. Kurushima, J.D.; Lipinski, M.J.; Gandolfi, B.; Froenicke, L.; Grahn, J.C.; Grahn, R.A.; Lyons, L.A. Variation of cats under domestication: genetic assignment of domestic cats to breeds and worldwide random-bred populations. Anim Genet 2013, 44, 311–324, doi:10.1111/age.12008.

6. Online Mendelian Inheritance in Animals, OMIA., Availabe online: https://omia.org/ (accessed on 28 Apr 2017).

7. Lenffer, J.; Nicholas, F.W.; Castle, K.; Rao, A.; Gregory, S.; Poidinger, M.; Mailman, M.D. OMIA (Online Mendelian Inheritance in Animals): an enhanced platform and integration into the Entrez search interface at NCBI. Nucleic Acids Res 2006, 34, D599–D601, doi:10.1093/nar/gkj152.

8. Gandolfi, B.; Alhaddad, H.; Affolter, V.K.; Brockman, J.; Haggstrom, J.; Joslin, S.E.K.; Koehne, A.L.; Mullikin, J.C.; Outerbridge, C.A.; Warren, W.C., et al. To the root of the curl: a signature of a recent selective sweep identifies a mutation that defines the Cornish rex cat breed. PLoS One 2013, 8, doi:10.1371/journal.pone.0067105.

9. Gandolfi, B.; Outerbridge, C.A.; Beresford, L.G.; Myers, J.A.; Pimentel, M.; Alhaddad, H.; Grahn, J.C.; Grahn, R.A.; Lyons, L.A., The naked truth: sphynx and Devon rex cat breed mutations in *KRT71*. Mamm Genome 2010, 21, 509–515, doi:10.1007/s00335-010-9290-6.

10. Filler, S.; Alhaddad, H.; Gandolfi, B.; Kurushima, J.D.; Cortes, A.; Veit, C.; Lyons, L.A.; Brem, G., Selkirk rex: morphological and genetic characterization of a new cat breed. J Hered 2012, 103, 727–733, doi:10.1093/jhered/ess039.

11. Gandolfi, B.; Alhaddad, H.; Joslin, S.E.K.; Khan, R.; Filler, S.; Brem, G.; Lyons, L.A. splice variant in *KRT71* is associated with curly coat phenotype of Selkirk rex cats. Sci Rep 2013, 3, doi:10.1038/Srep02000.

12. Irvine, A.D.; McLean, W.H. Human keratin diseases: the increasing spectrum of disease and subtlety of the phenotype-genotype correlation. Br J Dermatol 1999, 140, 815–828, doi:10.1046/j.1365-2133.1999.02810.x.

13. Smith, F.J.D. The molecular genetics of keratin disorders. Am J Clin Dermatol 2003, 4, 347–364, doi:10.2165/00128071-200304050-00005.

14. Visinoni, A.F.; Lisboa-Costa, T.; Pagnan, N.A.B.; Chautard-Freire-Maia, E.A. Ectodermal dysplasias: clinical and molecular review. Am J Med Genet 2009, *149A*, 1980–2002, doi:10.1002/ajmg.a.32864.

15. Wright, J.T.; Fete, M.; Schenider, H.; Zinser, M.; Koster, M.I.; Clarke, A.J.; Hadj-Rabia, S.; Tadini, G.; Pagnan, N.; Visinoni, A.F., et al. Ectodermal dysplasias: classification and organization by phenotype, genotype and molecular pathway. Am J Med Genet 2019, 179, 442–447, doi:10.1002/ajmg.a.61045.

16. Ahmad, W.; Ratterree, M.S.; Panteleyev, A.A.; Aita, V.M.; Sundberg, J.P.; Christiano, A.M. Atrichia with papular lesions resulting from mutations in the rhesus macaque (*Macaca mulattta*) hairless gene. Lab Anim 2002, 36, 61–67, doi:10.1258/0023677021911777.

17. Finocchiaro, R.; Portolan, B.; Damiani, G.; Caroli, A.; Budelli, E.; Bolla, P.; Pagnacco, G. The hairless (*hr*) gene is involved in the congenital hypotrichosis of Valle del Belice sheep. Genet Sel Evol 2003, 35, S147–S156, doi:10.1051/gse:2003023.

18. Barlund, C.S.; Clark, E.G.; Leeb, T.; Drogemuller, C.; Palmer, C.W. Congenital hypotrichosis and partial anodontia in a crossbreed beef calf. Can Vet J 2007, 48, 612–614.

19. Drogemuller, C.; Karlsson, E.K.; Hytonen, M.K.; Perloski, M.; Dolf, G.; Saino, K.; Lohi, H.; Lindblad-Toh, K.; Leeb, T. A mutation in hairless dogs implicates *FOXI3* in ectodermal development. Science 2008, 321, 1462, doi:10.1126/science.1162525.

20. Chen, Z.; Wang, Z.; Xu, S.; K., Z.; G., Y. Characterization of hairless (*Hr*) and *FGF5* genes provides insights into the molecular basis of hair loss in cetaceans. BMC Evol Biol 2013, 13, 34, doi:10.1186/1471-2148-13-34.

21. Parker, H.G.; Harris, A.; Dreger, D.L.; Davis, B.W.; Ostrander, E.A. The bald and the beautiful: hairlessness in domestic dog breeds. Phil Trans R Soc B 2017, 372, 20150488, doi:10.1098/rstb.2015.0488.

22. Hadji-Rasouliha, S.; Bauer, A.; Dettwiler, M.; Welle, M.M.; Leeb, T. A frameshift variant in the *EDA* gene in dachshunds with X-linked hypohidrotic ectodermal dysplasia. Anim Genet 2018, 49, 651–654, doi:10.1111/age.12729.

23. Abitbol, M.; Bosse, P.; Thomas, A.; Tiret, L. A deletion in *FOXN1* is associated with a syndrome characterized by congenital hypotrichosis and short life expectancy in Birman cats. PLoS One 2015, 10, e0120668, doi:10.1371/journal.pone.0120668.

24. David, V.A.; Menotti-Raymond, M.; Wallace, A.C.; Roelke, M.; Leighty, R.; Eizirik, E.; Hannah, S.S.; Nelson, G.; Schäffer, A.A.; Connelly, C.J., et al. Endongenous retrovirus insertion in the *KIT* oncogene determines white and white spotting in domestic cats. G3 2014, 4, 1881–1891, doi:10.1534/g3.114.013425.

25. Lyons, L.A.; Imes, D.I.; Rah, H.C.; Grahn, R.A. Tyrosinase mutations associated with Siamese and Burmese patterns in the domestic cat (*Felis catus*). Anim Genet 2005, 36, 119–126, doi:10.1111/j.1365-2052.2005.01253.x.

26. Imes, D.I.; Geary, L.A.; Grahn, R.A.; Lyons, L.A., Albinism in the domestic cat (*Felis catus*) is associated with a tyrosinase (*TYR*) mutation. Anim Genet 2006, 37, 175–180, doi:10.1111/j.1365-2052.2005.01409.x.

27. Creel, D.; Hendrickson, A.E.; Leventhal, A.G. Retinal projections in tyrosinase-negative albino cats. J Neurosci 1982, 2, 907–911, doi:10.1523/JNEUROSCI.02-07-00907.1982.

28. LeRoy, M.L.; Senter, D.A.; Kim, D.Y.; Gandolfi, B.; Middleton, J.R.; Bouhan, D.M.; Lyons, L.A. Clinical and histologic description of lykoi cat hair coat and skin. Jap J Vet Dermatol 2016, 22, 179–191.

29. TICA. The International Cat Association. Availabe online: (accessed on 01 May 2020).

30. Lyons, L.A.; Creighton, E.K.; Alhaddad, H.; Beale, H.C.; Grahn, R.A.; Rah, H.; Maggs, D.J.; Helps, C.R.; Gandolfi, B. Whole genome sequencing in cats, identifies new models for blindness in *AIPL1* and somite segmentation in *HES7*. BMC Genomics 2016, 17, 265, doi:10.1186/s12864-016-2595-4.

31. Mauler, D.A.; Gandolfi, B.; Rineiro, C.R.; O’Brien, D.P.; Spooner, J.L.; Lyons, L.A. Precision medicine in cats: novel Niemann-Pick type C1 diagnosed by whole-genome sequencing. J Vet Intern Med 2017, 31, 539–544, doi:10.1111/jvim.14599.

32. Oh, A.; Pearce, J.W.; Gandolfi, B.; Creighton, E.K.; Suedmeyer, W.K.; Michael Selig, M.; Bosiack, A.P.; Castaner, L.J.; Whiting, R.E.,H.; Belknap, E.B., et al. Early-Onset Progressive Retinal Atrophy Associated with an *IQCB1* Variant in African Black-Footed Cats (*Felis nigripes*). Sci Rep 2017, 10.1038/srep43918, doi:10.1038/srep43918.

33. Buckley, R.M.; Grahn, R.A.; Gandolfi, B.; Herrick, J.R.; Kittleson, M.D.; Bateman, H.L.; Newsom, J.; Swanson, W.F.; Prieur, D.J.; Lyons, L.A. Assisted reproduction mediated resurrection of a feline model for Chediak-Higashi syndrome caused by a large duplication in *LYST*. Sci Rep 2020, 10, 64, doi:10.1038/s41598-019-56896-9.

34. Sambrook, J.; Russell, D.W. Preparation and analysis of eukaryotic genomic DNA. In Molecular cloning: a laboratory manual, 3rd ed.; Sambrook, J., Russell, D.W., Eds. Cold Spring Laboratory Press: Cold Spring Harbor, New York, USA, 2001; pp. 6.11–16.14.

35. Lipinski, M.J.; Amigues, Y.; Blasi, M.; Broad, T.E.; Cherbonnel, C.; Cho, G.J.; Corley, S.; Daftari, P.; Delattre, D.R.; Dileanis, S., et al. An international parentage and identification panel for the domestic cat (*Felis catus*). Anim Genet 2007, 38, 371–377, doi:10.1111/j.1365-2052.2007.01632.x.

36. Toonen, R.J.; Hughes, S. Increased throughput for fragment analysis on an ABI Prism 377 automated sequencer using a membrane comb and STRand software. BioTechniques 2001, 31, 1320–1324.

37. Buckley, R.M.; Davis, B.W.; Brashear, W.A.; Farias, F.H.; Kuroki, K.; Graves, T.; Hillier, L.W.; Kremitzki, M.; Li, G.; Middleton, R. A new domestic cat genome assembly based on long sequence reads empowers feline genomic medicine and identifies a novel gene for dwarfism. bioRxiv 2020, 896258.

38. Li, H. Aligning sequence reads, clone sequences and assembly contigs with BWA-MEM. arXiv preprint arXiv:1303.3997. 2013.

39. McKenna, A.; Hanna, M.; Banks, E.; Sivachenko, A.; Cibulskis, K.; Kernytsky, A.; Garimella, K.; Altshuler, D.; Gabriel, S.; Daly, M., et al. The Genome Analysis Toolkit: a MapReduce framework for analyzing next-generation DNA sequencing data. Genome Res 2010, 20, 1297–1303, doi:10.1101/gr.107524.110.

40. Poplin, R.; Ruano-Rubio, V.; DePristo, M.; Fennell, T.; Carneiro, M.; Van der Auwera, G.; Kling, D.; Gauthier, L.; Levy-Moonshine, A.; Roazen, D. Scaling accurate genetic variant discovery to tens of thousands of samples. bioRxiv 2017, 201178.

41. Li, H.; Handsaker, B.; Wysoker, A.; Fennell, T.; Ruan, J.; Homer, N.; Marth, G.; Abecasis, G.; Durbin, R.; Genome Project Data Processing, S. The Sequence Alignment/Map format and SAMtools. Bioinformatics 2009, 25, 2078–2079, doi:10.1093/bioinformatics/btp352.

42. Cunningham, F.; Achuthan, P.; Akanni, W.; Allen, J.; Amode, M.R.; Armean, I.M.; Bennett, R.; Bhai, J.; Billis, K.; Boddu, S., et al. Ensembl 2019. Nucleic Acids Res 2019, 47, D745–D751, doi:10.1093/nar/gky1113.

43. Gandolfi, B.; Daniel, R.J.; O’Brien, D.P.; Guo, L.T.; Youngs, M.D.; Leach, S.B.; Jones, B.R.; Shelton, G.D.; Lyons, L.A. A novel mutation in *CLCN1* associated with feline myotonia congenita. PLoS One 2014, 9, e109926, doi:10.1371/journal.pone.0109926.

44. Cachon-Gonzalez, M.B.; Fenner, S.; Coffin, J.M.; Moran, C.; Best, S.; Stoye, J.P. Structure and expression of the hairless gene of mice. Proc Natl Acad Sci USA 1994, 91, 7717–7721, doi:10.1073/pnas.91.16.7717.

45. Ahmad, W.; Faiyaz ul Haque, M.; Brancolini, V.; Tsou, H.C.; ul Haque, S.; Lam, H.; Aita, V.M.; Owen, J.; deBlaquiere, M.; Frank, J., et al. Alopecia universalis associated with a mutation in the human hairless gene. Science 1998, 279, 720–724, doi:10.1126/science.279.5351.720.

46. Menotti-Raymond, M.; David, V.A.; Weir, B.S.; O’Brien, S.J. A population genetic database of cat breeds developed in coordination with a domestic cat STR multiplex. J Forensic Sci 2012, 57, 596–601, doi:10.1111/j.1556-4029.2011.02040.x.

47. Robinson, R. Devon rex-a third rexoid coat mutation in the cat. Genetica 1969, 40, 597–599, doi:10.1007/BF01787284.

48. Searle, A.G.; Jude, A.C. The ‘rex’ type coat in the domestic cat. J Genet 1956, 506–512.

49. Bult, C.J.; Eppig, J.T.; Kadin, J.A.; Richardson, J.E.; Blake, J.A.; Group, M.G.D. The mouse genome database (MGD): mouse biology and model systems. Nucleic Acids Res 2008, 36, D724–D728, doi:10.1093/nar/gkm961.

50. Brooke, H.C. Hairless mice. J Hered 1926, 17, 173–174, doi:10.1093/oxfordjournals.jhered.a102700.

51. Ahmad, W.; Panteleyev, A.A.; Christiano, A.M. The molecular basis of congential atrichia in humans and mice: mutations in the hairless gene. J Investig Dermatol Symp Proc 1999, 4, 240–243, doi:10.1038/sj.jidsp.5640220.

52. Klein, I.; Bergman, R.; Indelman, M.; Sprecher, E. A novel missense mutation affecting the human hairless thyroid receptor interacting domain 2 causes congenital atrichia. J Investig Dermatol 2002, 119, 920–922, doi:10.1046/j.1523-1747.2002.00268.x.

53. Moraitis, A.N.; Giguere, V. The co-repressor hairless protects ROR-alpha orphan nuclear receptor from proteasome mediated degradation. J Biol Chem 2003, 278, 52511–52518, doi:10.1074/jbc.M308152200.

54. Hsieh, J.-C.; Sisk, J.M.; Jurutka, P.W.; Haussler, C.A.; Slater, S.A.; Haussler, M.R.; Thompson, C.C. Physical and functional interaction between the vitamin D receptor and hairless corepressor, two proteins required for hair cycling. J Biol Chem 2003, 278, 38665–38674, doi:10.1074/jbc.M304886200.

55. Potter, G.B.; Beaudoin III, G.M.J.; DeRenzo, C.L.; Zarach, J.M.; Chen, S.H.; Thompson, C.C. The *hairless* gene mutated in congenital hair loss disorders encodes a novel nuclear receptor corepressor. Genes Dev 2001, 15, 2687–2701, doi:10.1101/gad.916701.

56. Wen, Y.; Liu, Y.; Xu, Y.; Zhou, Y.; Hua, R.; Wang, K.; Sun, M.; Li, Y.; Yang, S.; Zhang, X.-J., et al. Loss-of-function mutations of an inhibitory upstream ORF in the human hairless transcript cause Marie Unna hereditary hypotrichosis. Nat Genet 2009, 41, 228–233, doi:10.1038/ng.276.

57. Landrum, M.J.; Lee, J.M.; Benson, M.; Brown, G.R.; Chao, C.; Chitipiralla, S.; Gu, B.; Hart, J.; Hoffman, D.; Jang, W., et al. ClinVar: improving access to variant interpretations and supporting evidence. Nucleic Acids Res 2018, 46, D1062–D1076, doi:10.1093/nar/gkx1153.

58. Aita, V.M.; Ahmad, W.; Panteleyev, A.A.; Kozlowska, U.; Kozlowska, A.; Gilliam, T.C.; Jablonska, S.; Christiano, A.M., A novel missense mutation (C622G) in the zinc finger domain of the human hairless gene associated with congenital atrichia with papular lesions. Experim Dermatol 2001, 9, 157–162, doi:10.1034/j.1600-0625.2000.009002157.x.

59. Hytonen, M.K.; Lohi, H. A frameshift insertion in *SGK3* leads to recessive hairlessness in Scottish Deerhounds: a candidate gene for human alopecia conditions. Human Genet 2019, 138, 535–539, doi:10.1007/s00439-019-02005-9.

60. Parker, H.G.; Whitaker, D.T.; Harris, A.C.; Ostrander, E.A. Whole genome analysis of a single Scottish deerhound family provides independent corroboration that a *SGK3* coding variant leads to hairlessness. G3 2020, 10, 293–297, doi:10.1534/g3.119.400885.

61. Cunningham, F.; Achuthan, P.; Akanni, W.; Allen, J.; Amode, M R.; Armean, I.M.; Bennett, R.; Bhai, J.; Billis, K.; Boddu, S., et al. Ensembl 2019. Nucleic Acids Res 2018, 47, D745–D751, doi:10.1093/nar/gky1113.

62. Drogemuller, C.; Rufenacht, S.; Wichert, B.; Leeb, T. Mutations within the *FGF5* gene are associated with hair length in cats. Anim Genet 2007, 38, 218–221, doi:10.1111/j.1365-2052.2007.01590.x.

63. Kehler, J.S.; David, V.A.; Schaffer, A.A.; Bajema, K.; Eizirik, E.; Ryugo, D.K.; Hannah, S.S.; O’Brien, S.J.; Menotti-Raymond, M., Four independent mutations in the feline Fibroblast Growth Factor 5 gene determine the long-haired phenotype in domestic cats. J Hered 2007, 98, 555–566, doi:10.1093/jhered/esm072.

64. Buckingham, K.J.; McMillin, M.J.; Brassil, M.M.; Shively, K.M.; Magnaye, K.M.; Cortes, A.; Weinmann, A.S.; Lyons, L.A.; Bamshad, M.J. Multiple mutant *T* alleles cause haploinsufficiency of brachyury and short tails in Manx cats. Mamm Genome 2013, 10.1007/s00335-013-9471-1, doi:10.1007/s00335-013-9471-1.

65. Kaelin, C.B.; Xu, X.; Hong, L.Z.; David, V.A.; McGowan, K.A.; Schmidt-Kuntzel, A.; Roelke, M.E.; Pino, J.; Pontius, J.; Cooper, G.M., et al. Specifying and sustaining pigmentation patterns in domestic and wild cats. Science 2012, 337, 1536–1541, doi:10.1126/science.1220893.

66. Gandolfi, B.; Alamri, S.; Darby, W.G.; Adhikari, B.; Lattimer, J.C.; Malik, R.; Wade, C.M.; Lyons, L.A.; Cheng, J.; Bateman, J.F., et al. A dominant *TRPV4* variant underlies osteochondrodysplasia in Scottish fold cats. Osteoarthritis Cartilage 2016, 24, 1441–1450, doi:10.1016/j.joca.2016.03.019.

67. Menotti-Raymond, M.; David, V.A.; Stephens, J.C.; Lyons, L.A., O’Brien, S.J. Genetic individualization of domestic cats using feline STR loci for forensic applications. J Forensic Sci 1997, 42, 1039–1051, doi:10.1520/JFS14258J.

68. Menotti-Raymond, M.; David, V.A.; Lyons, L.A.; Schaffer, A.A.; Tomlin, J.F.; Hutton, M.K.; O’Brien, S.J. A genetic linkage map of microsatellites in the domestic cat (*Felis catus*). Genomics 1999, 57, 9–23, doi:10.1006/geno.1999.5743.

